# Rapid bacteria-phage coevolution drives the emergence of multi-scale networks

**DOI:** 10.1101/2023.04.13.536812

**Authors:** Joshua M. Borin, Justin J. Lee, Adriana Lucia-Sanz, Krista R. Gerbino, Joshua S. Weitz, Justin R. Meyer

**Affiliations:** Division of Biological Sciences, University of California San Diego, San Diego, CA, USA; School of Physics, Georgia Institute of Technology, Atlanta, GA, USA; School of Biological Sciences, Georgia Institute of Technology, Atlanta, GA, USA; Institut de Biologie, École Normale Supérieure, Paris, France

## Abstract

Interactions between species have catalyzed the evolution of multiscale ecological networks–including both nested and modular elements that regulate the function of diverse communities. One common assumption is that such complex pattern formation requires long evolutionary timescales, spatial isolation, or other exogenous processes. Here we show that multiscale network structure can evolve rapidly under simple ecological conditions without spatial structure. In just 21 days of laboratory coevolution, *Escherichia coli* and bacteriophage Φ21 coevolve and diversify to form elaborate cross-infection networks. By measuring ∼10,000 phage–bacteria infections and testing the genetic basis of interactions, we identify the mechanisms that create each component of the multiscale pattern. Initially, nested patterns form through an arms race where hosts successively lose the original receptor (LamB) and phages evolve to use a second (OmpC) and then a third (OmpF) receptor. Next, modules form when the cost of losing the third receptor, OmpF, increases and bacteria evolve resistance mutations that modify the OmpF receptors’ extramembrane loops. In turn, phages evolve adaptations that facilitate specialized interactions with different OmpF variants. Nestedness reemerges within modules as bacteria evolve increased resistance and phages enhance infectivity against module-specific receptor variants. Our results demonstrate how multiscale networks evolve in parasite-host systems, illustrating Darwin’s idea that simple adaptive processes can generate *entangled banks* of ecological interactions.

---

Recent analyses of ecological networks reveal structural patterns that recur in disparate ecosystems^1–6^. Two commonly observed patterns include nestedness, where specialized interactions are hierarchically embedded within generalized interactions^1,7^, and modularity, where specialized interactions form within—but not between—groups, generating distinct “modules” ^2,6^. Multiscale networks are also found in large-scale analyses of natural community interactions, whereby modularity is observed at broad scales and nestedness is observed within modules^3–5,8,9^. How multiscale patterns emerge within ecological networks is an open question. It has been suggested that formation of such intricate structures requires geographic separation^8,10,11^, long evolutionary timescales^12,13^, or other external drivers^5,14^. However, it has been hypothesized that coevolution between interacting species could lead to such complexity even on short time scales and in simple, closed communities^15^.

Bacteria and their viruses (phages) provide tractable systems for studying how complex ecological networks evolve^4,16,17^. By isolating phages and bacteria from a given ecosystem and measuring all-by-all pairwise infections, it is possible to construct phage-bacteria interaction networks (PBINs) and analyze their underlying structure. Under controlled laboratory settings, phage-bacteria coevolution experiments rapidly generate rich ecological networks, allowing researchers to study the mechanisms underlying their formation^7,18–20^. For example, arms race dynamics produce nested patterns when bacteria evolve escalating resistance and phages counter with increasing infectivity^4,7,21^. Modules can form due to fluctuating selection dynamics^18,20,22^ or the presence of different phage defense elements that confer specialized immunity^23^. However, experiments have been unable to reproduce the complex multiscale PBINs observed in nature without artificially providing spatial isolation between evolving subpopulations^24^, reaffirming the common assumption that spatial structure and/or other external influences are a prerequisite for the evolution of intricate ecological networks. Here we decipher the molecular mechanisms of parasite-host coevolution and demonstrate that the process of coevolution itself is sufficient to drive the rapid emergence of complex multiscale networks in simple ecological contexts.

## Coevolution drives multiscale networks

We began by inoculating 3 well-mixed cultures with isogenic populations of *Escherichia coli* K-12^25^ and a lytic strain of bacteriophage 21 (Φ21)^26^. Cultures were incubated at 37°C and every 24 hours, 1% of each community was propagated into new media, allowing bacteria and phage to coevolve for 21 days. Each day, we tested phage populations for receptor use expansion and found that in 2 of 3 replicates, phages evolved to use two new receptors (described below). As a preliminary test, we then isolated a single phage from each population and timepoint. By comparing phage receptor usage in population 2, we discovered that phages initially expanded and then contracted their receptor use, consistent with an initial arms race transitioning to more specialized interactions. Therefore, we focused our efforts on population 2 (replicate experiments demonstrated that this transition was repeatable, a topic that we revisit later in the manuscript).

To analyze the dynamics in population 2, we isolated coevolved phages (n=74) and bacteria (n=128) from communities every third day and measured all 9,472 pairwise infections to generate a PBIN. Initially, the network showed a nested pattern, as bacteria and phages evolved broader resistance and host range over time (Fig. 1a; Extended Data Fig. 1, p<0.001 for bacteria and phage, linear model). However, the nested pattern was disrupted between days 18 and 21; phages isolated on day 21 broadened their host range to infect day-18 hosts while losing the ability to infect day-15 hosts. This tit-for-tat pattern led us to hypothesize that specialized interactions may have formed, creating modules in the network.

**Fig. 1.**
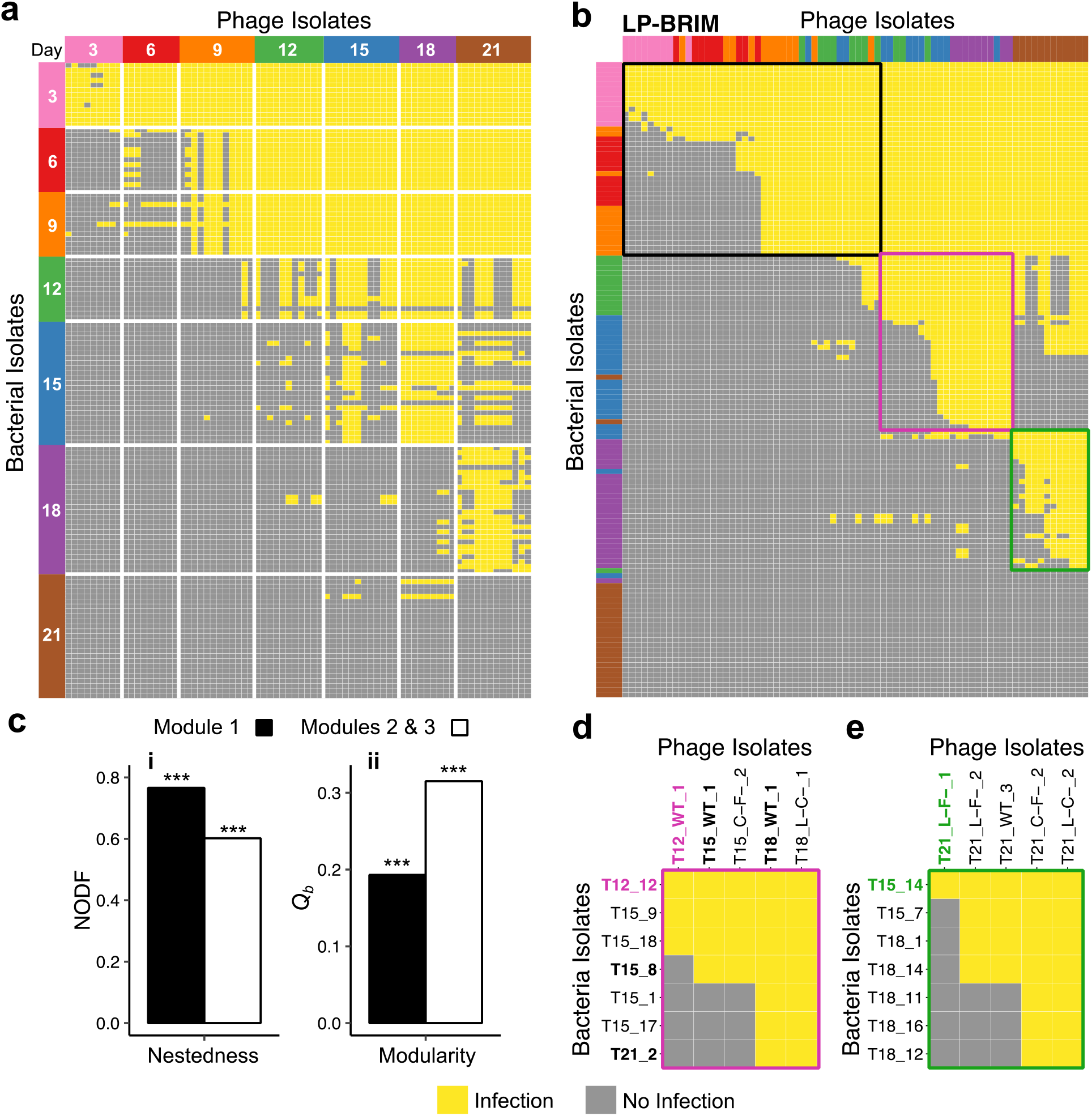
Phage–bacteria coevolution generates multiscale network patterns in 21 days. **a**, Phage– bacteria interaction network (PBIN) comprised of 9,472 pairwise infections between 128 *E. coli* and 74 Φ21 strains isolated from various days of coevolution. **b**, Community detection of the PBIN using the LP-BRIM algorithm reveals 3 modules (1, 2, and 3 indicated in black, pink, green, respectively: *Q*_*b*_=0.2100***, N=3). **c**, Nestedness (NODF) and modularity (LP-BRIM) of interactions between isolates from early (**i**, days 3-9: NODF=0.7655, *Q*_*b*_=0.1935, N=2) and late (**ii**, days 12-21: NODF=0.6019, *Q*_*b*_=0.3151, N=2) in the coevolution (p<0.0001 ***). **d** and **e**, Infections between representative isolates from modules 2 and 3 were remeasured and recapitulate within-module nestedness. **d** and **e** serve as roadmaps for proceeding figures. Label aesthetics are applied consistently for continuity.

We tested for the emergence of modules using the LP-BRIM (Label Propagation and Bipartite Recursively Induced Modules) algorithm which searches for community configuration in a bipartite network that maximizes the modularity metric (0 < *Q*_*b*_ ≤ 1, where 1 denotes fully modular; see Methods)^27,28^. The LP-BRIM algorithm identified 3 modules (Fig. 1b; *Q*_*b*_=0.210, p<0.0001; Extended Data Fig. 2b). The first module includes bacteria isolates from days 3-9 and (nearly exclusively) early phage isolates from days 3-12. In contrast, modules 2 and 3 include phage and bacteria isolates exclusively from days 12-21. Next, we tested all 3 modules for nestedness, using the “Nestedness metric based on Overlap and Decreasing Fill” (NODF) method^29^ (see Methods); all 3 modules were significantly nested (module 1 NODF=0.766, p<0.0001; module 2 NODF=0.802, p<0.0001; module 3 NODF= 0.777, p<0.0001). Notably, phage-bacteria infections amongst early isolates in module 1 were more nested (NODF=0.766, p<0.0001) than later isolates in modules 2 and 3 combined (NODF=0.602, p<0.0001) while infection between later isolates were significantly more modular (*Q*_*b*_=0.315, p<0.0001) than early isolates (*Q*_*b*_=0.194, p<0.0001)–a result of reduced cross-infections between phages of module 2 or 3 with bacteria of module 3 or 2, respectively (Fig. 1c).

To validate the nested pattern within modules, as well as the repeatability of PBIN measurements, we remeasured representative interactions within modules 2 and 3, which recapitulated nestedness within each module (Fig. 1d, e) and we independently remeasured 3654-interaction subsets of the full PBIN twice, which were significantly correlated with Fig. 1a (p<0.001, Mantel test, Extended Data Fig. 3). After demonstrating that coevolution between *E. coli* and phage Φ21 led to the emergence of a multiscale, nested-modular network, we endeavored to determine the evolutionary and molecular processes responsible for each pattern: 1) initial nestedness, 2) the formation of modules, and 3) nestedness reemerging within modules.

### Initial nestedness

We first investigated the factors driving the emergence of nestedness in module 1. Prior studies of coevolution between *E. coli* and phage λ show that arms race dynamics produce nested patterns as bacteria evolve resistance by mutating λ-receptor LamB and phages counter by innovating to use a new receptor, OmpF^21,31,32^. We tested whether similar arms race dynamics drove nestedness in module 1 by measuring the frequency of host receptor mutations and phage receptor use in the population from days 0–12 (Fig. 2a). Population dynamics revealed multiple cycles of an arms race, as bacteria sequentially mutated outer membrane proteins LamB, OmpC, and OmpF, and phages that initially infected through LamB innovated to use OmpC and then OmpF. We then determined the receptor use of individual phage isolates by spotting them onto different lawns of bacteria with 2 of 3 relevant receptors deleted. This revealed that phages from day 3 could use either LamB or OmpC alone as receptors and phages from day 9 could use LamB, OmpC, or OmpF, representing the first instance of a documented triple-receptor phage we are aware of (Fig. 2b). Lastly, to confirm that consecutive receptor mutations and innovations were responsible for nestedness in module 1, we compared and found direct concordance between phage isolates from days 0, 3, and 9 infecting naturally coevolved bacteria and hosts genetically engineered with nonsense or deletion mutations in corresponding receptors (Fig. 2c). Altogether, our results confirm that multiple rounds of host receptor loss and phage receptor-use gain drove initial nestedness in the network.

**Fig. 2.**
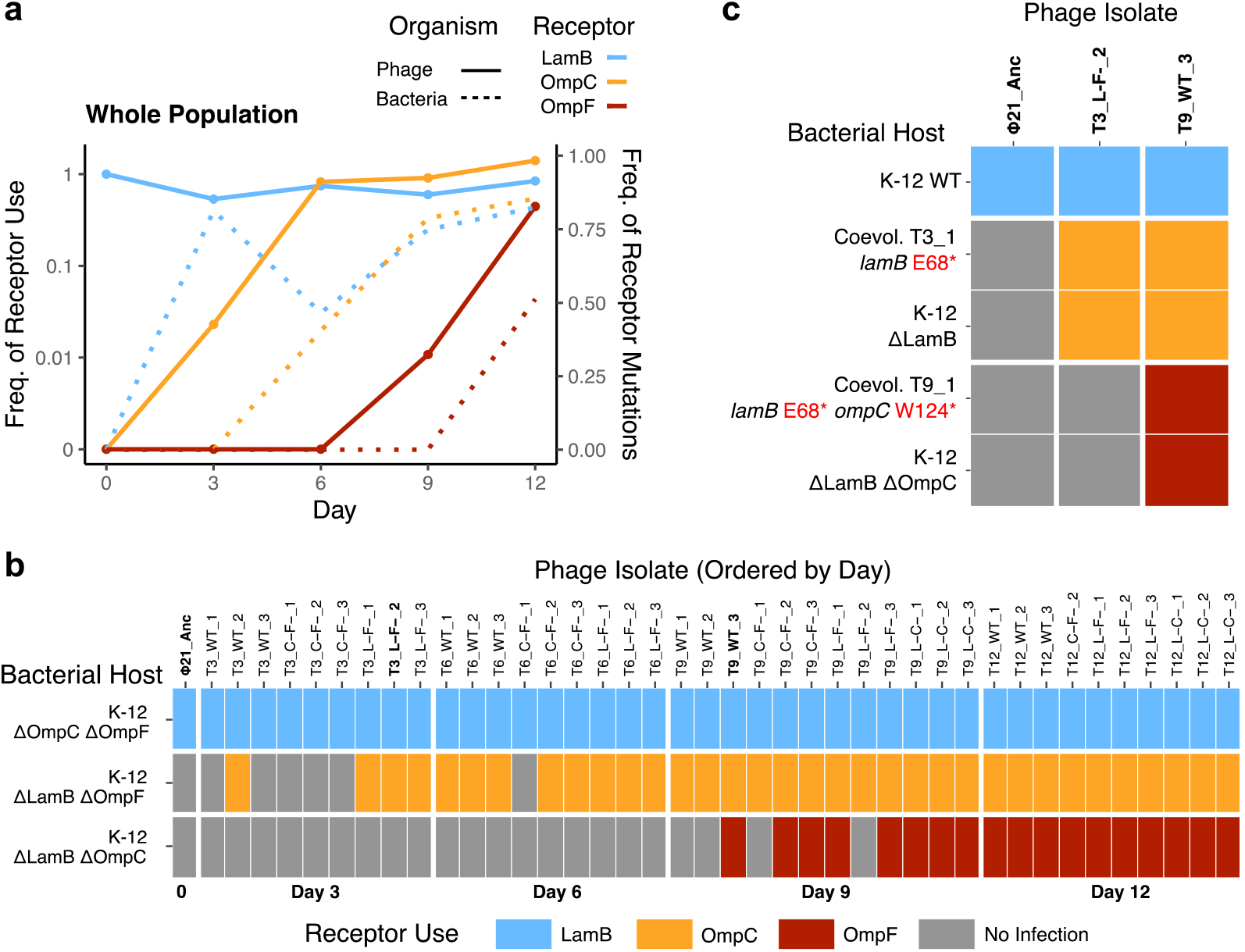
Initial nestedness is driven by multiple cycles of host receptor loss and phage receptor innovation. **a**, Frequency of phage receptor use (solid) and bacterial receptor mutations (dotted) in populations across coevolution days 0–12. Frequency of receptor use was calculated as the titer of phages on ΔOmpC ΔOmpF, ΔLamB ΔOmpF, or ΔLamB ΔOmpC K-12 hosts relative to K-12 WT. Frequency of receptor mutations was calculated as the frequency of whole population sequencing reads with mutations affecting each receptor. **b**, Ability of 41 phage isolates from coevolution days 0–12 to use host receptors LamB, OmpC, or OmpF, determined by spotting phages on agar infused with different dual-receptor knockout hosts. **c**, Ability of phage isolates with expanding receptor use to infect naturally coevolved and genetically engineered bacteria with mutations successively eliminating receptors.

By day 9, evolved bacteria had lost the first two receptors (LamB and OmpC) and phages countered by evolving to use a third receptor, OmpF. We discovered that bacteria could evolve complete resistance by deleting the final OmpF receptor (day 12, 1 of 13 isolates), yet completely resistant mutants did not increase in frequency until day 21 (23 of 25 isolates). Bacterial fitness competitions revealed that the sequential loss of LamB, OmpC, and then OmpF receptors correlated with 14%, 31%, and 50% reductions in growth rate, respectively, with completely resistant isolates paying immense costs (Extended Data Fig. 4). These results support a coevolutionary mechanism by which bacteria maintained the OmpF receptor until phage-induced lysis became sufficiently intense to select for complete resistance between days 18 and 21.

### Module formation

The separation of modules 2 and 3, comprised of phages and bacteria isolated on Day 12 and beyond implies that as phages gained the ability to infect new hosts, the phages lost infectivity on other, evolved hosts (see Fig. 1b, d, e, Extended Data Fig. 2). Because bacteria retained the OmpF receptor and began evolving nucleotide substitutions within *ompF*, we hypothesized that modules formed due to specialized interactions between different phages and variants of OmpF. To investigate, we focused on bacteria and phages with the narrowest resistance or infectivity range within each module, as interactions between these isolates initiate the divergence of modules in the network. We refer to these strains as “founder” isolates. As expected, founder phages could infect within, but not between, modules 2 and 3 (Fig. 3a). Importantly, this tit-for-tat pattern was recapitulated when phages were tested against ΔLamB ΔOmpC hosts engineered with *ompF* mutations from founder bacteria, confirming that interactions between different phages and OmpF variants drove module formation (Fig. 3a).

**Fig. 3.**
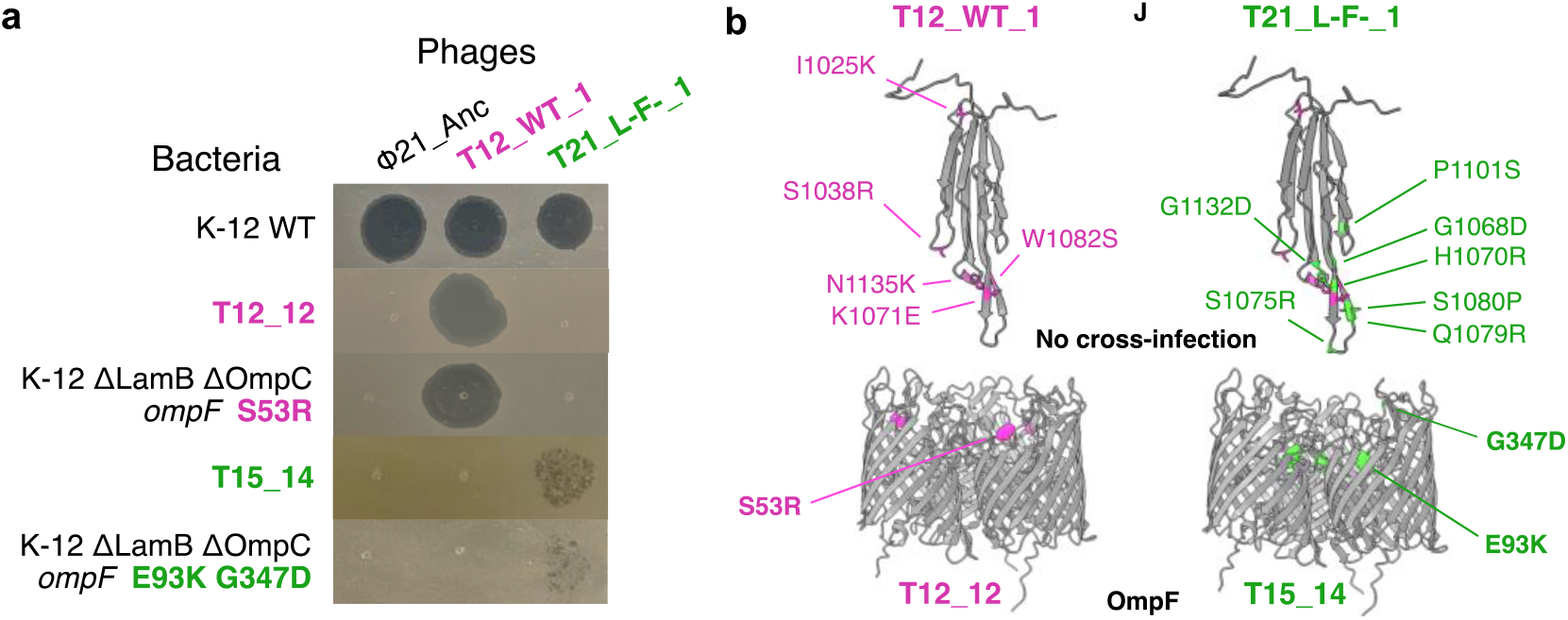
Modules form due to specialized interactions between different phages and OmpF variants. **a**, Infection assay of Φ21 and module founder phages spotted onto lawns of K-12 WT, module founder bacteria, and ΔLamB ΔOmpC hosts engineered with *ompF* mutations from corresponding founder bacteria. **b**, AlphaFold predicted structures of the phage host-recognition central tail fiber protein J (top) and solved structure of host OmpF receptor (bottom). Mutations in module 1 and 2 isolates are annotated in pink and green, respectively. Phage T21_L-F-_1 has 7 mutations (green) in addition to the 5 mutations present in phage T12_WT_1 (pink). OmpF mutants do not share any mutations.

To illustrate this result, we mapped bacterial mutations onto solved structures of OmpF and phage mutations onto predicted structures of the host-recognition protein, (similar to λ central tail fiber protein, J) (Fig. 3b). We found that OmpF mutations were located on extramembrane loops and J mutations were concentrated on extruding finger-like projections. The exposure of these sites on the outer surface of the host receptor and phage tail fiber strongly suggests that infectivity between founder phages and bacteria is governed by specific binding between accessible surfaces on variants of J and OmpF. Interestingly, although infectivity between phages and OmpF variants is modular, the phage phylogeny is nested. Module 3 phage T21_L-F-_1 is a descendant of module 2 phage T12_WT_1, indicating that 7 additional mutations caused it to gain function on module 3 OmpF variants and lose function on module 2 OmpF variants. Phage mutations with the capacity to cause simultaneous gain and loss of hosts have been observed previously^22,33^ and similar breakdowns between phylogenetic and phenotypic patterns have been observed in *E. coli–*λ coevolution^21^.

### Nestedness within modules

Lastly, we investigated the final pattern in our multiscale PBIN: the formation of nestedness within modules. In line with our observations for module formation, we hypothesized that additional *ompF* and *J* mutations produced nestedness within modules. Focusing on module 2, we used a similar approach as described above. First, we measured interactions between representative phages and bacteria, recapitulating nestedness within the module (Fig. 4a). Notably, phylogenies of phage and bacterial isolates were also nested. As phages accumulated *J* mutations and bacteria accumulated *ompF* mutations, they gained increasing host and resistance range, respectively, although different *ompF* mutations conferred different amounts of protection. When we tested the same phages on ΔLamB ΔOmpC hosts engineered with *ompF* mutations from coevolved bacteria, we found the same pattern, confirming that within-module nestedness was driven by the accumulation of mutations in *J* and *ompF* (Fig. 4b).

**Fig. 4.**
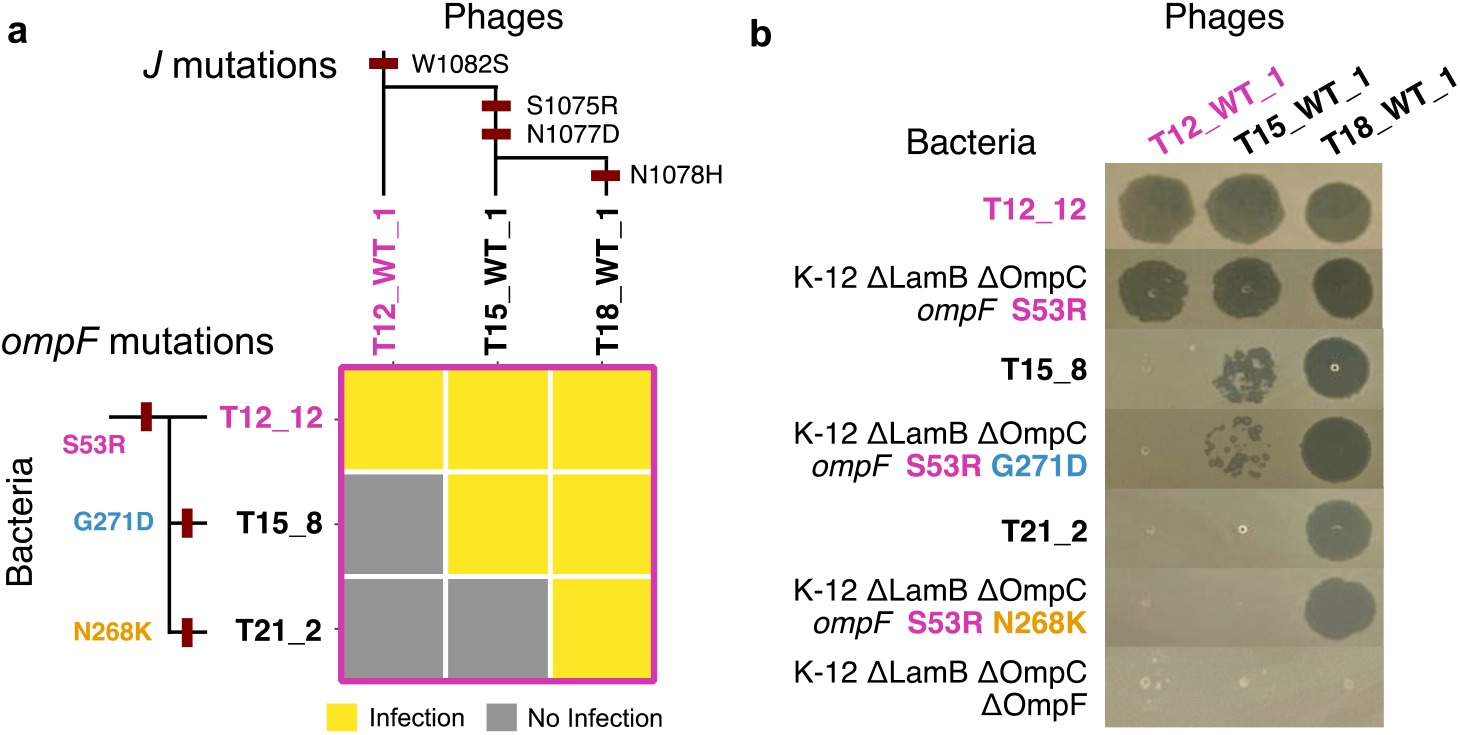
Lock-and-key arms race dynamics create within-module nestedness. **a**, Phylogenies of *J* and *ompF* mapped onto a PBIN of representative phage and bacterial isolates from module 2 (see Fig. 1d). Mutations are indicated on the phylogeny in red and OmpF mutations are annotated in pink, blue, and orange. **b**, Infection assay of phages spotted onto lawns of coevolved bacteria or ΔLamB ΔOmpC hosts engineered with *ompF* mutations from corresponding bacteria.

## Discussion

In this study, we demonstrated that coevolutionary processes are sufficient to generate complex multiscale ecological networks rapidly and without external influences or spatial structure. By determining the basis of key interactions, we revealed the evolutionary and molecular mechanisms underlying three major patterns in the network (summarized in Fig. 5): Initially, nestedness emerges as bacteria lose receptors and phages innovate to use new receptors through multiple cycles of an arms race. Modules form when bacteria are forced to retain the final OmpF receptor and phages evolve specialized interactions with mutated receptor variants. Finally, nestedness reemerges within modules as phages and bacteria accumulate mutations to increase infectivity and resistance via lock-and-key arms race dynamics.

**Fig. 5.**
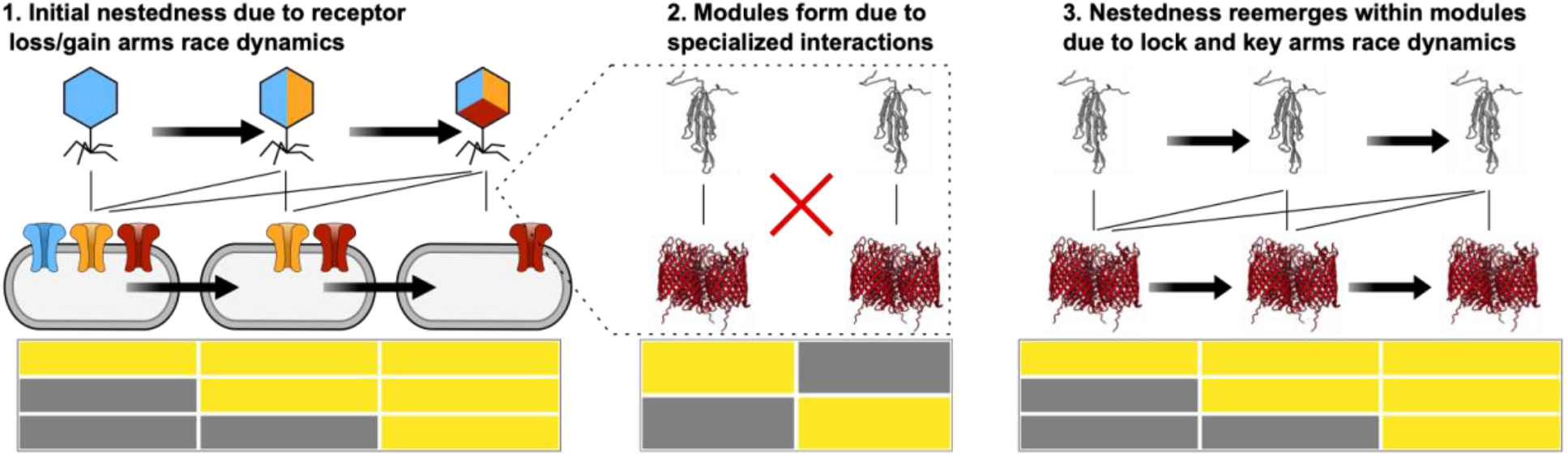
Mechanisms underlying three key patterns in the multiscale network. 1) Initial nestedness emerges due to multiple cycles of host receptor loss and phage receptor use gain. 2) Modules form due to specialized interactions between different phages and variants of the final receptor, OmpF. 3) Nestedness reemerges within modules due to lock-and-key arms race dynamics as mutations accumulate in phages and host receptors.

Notably, key features of the coevolution between *E. coli* and Φ21 were repeatable: Φ21 evolved to use a second receptor, OmpC in each of the 3 originally conducted replicates and 9 additional replicates conducted later in the same fashion. In 5 of 12 of these replicates, Φ21 evolved to use a third receptor, OmpF (Extended Data Fig. 5). In every case where OmpF use evolved, such use arose after phages evolved the ability to infect hosts via OmpC, suggesting similar coevolutionary mechanisms drove the dynamics.

Previous observations of geographically isolated modules have suggested that spatial separation assists in the formation of such patterns^5,8,9,24^. While it is reasonable to expect that space plays a role^34^, it is also possible that modules may separate in space after they form in sympatry. Similar pitfalls have been encountered when inferring past speciation processes from modern species distributions^35–37^. Instead, our finding provides a tractable experimental example of how diversity begets further diversity and, in turn, complex interaction structures^38,39^ akin to Darwin’s entangled bank^40^. While limits on diversity must exist, it seems that life–even in simple environments–is evolving far from these bounds.

## Acknowledgements

We thank Sherwood Casjens and the Ry Young Laboratory for providing bacteriophage 21 (Φ21) and its annotated genome. We thank Sarah Ardell, the Kryazhimskiy Lab, and other members of the Meyer Lab for useful discussions and comments on the manuscript. This work was supported by grants from the Howard Hughes Medical Institute Emerging Pathogens Initiative (7012574), National Institutes of Health (T32GM007240), the Army Research Office (W911NF1910384), and the Chaires Blaise Pascal program of the Île-de-France region.

## Author contributions

JMB and JRM conceptualized the study. JMB, JLL, and KRG performed the experiments. JMB and ALS conducted data analyses and made figures. JSW and JRM supervised the study. JMB and JRM wrote the manuscript. All authors edited the manuscript.

## Correspondence and requests for materials

should be addressed to JRM.

## Methods

### Strains

To study phage–bacteria coevolution, we used *Escherichia coli* strain K-12 BW25113 (WT)^25^ and bacteriophage 21 (Φ21, GenBank: OL657228)^26^. Φ21 is a “lambdoid” phage, sharing genetic similarity and a similar life cycle to phage λ, but its evolution is not well studied^41^. Using the wildtype, lysogenic Φ21, we inserted 2 stop mutations and a 1 bp frameshift deletion in the *cI*–like repressor gene (CPT_21_51) using techniques described below. We sequenced whole phage genomes which confirmed that *cI* engineering was successful and revealed one mutational difference between our Φ21 and the GenBank reference (G18903A, central tail fiber CPT_21_21, similar to phage λ gene *J*). To characterize phage receptor use and elucidate the molecular mechanisms underlying key interactions in our study, we also utilized a suite of hosts derived from the KEIO gene knockout collection^25^ with additional genetic mutations that functionally knock out 2 of the 3 relevant receptors. ΔLamB ΔOmpC and ΔOmpC ΔOmpF hosts were derived from KEIO strain JW2203 (*ΔompC*) and have nonsense mutations in *lamB* or *ompF*. ΔLamB ΔOmpF was derived from KEIO strain JW0912 (*ΔompF*) and has a nonsense mutation in *lamB*.

### Initial coevolution experiment

*E. coli* WT (∼10^5^ cells) and lytic Φ21 (∼10^5^ particles) were inoculated into three replicate 50-mL flasks containing 10 mL Tris-LB media (0.28 g K_2_HPO_4_, 0.08 g KH_2_PO_4_, 1 g (NH_4_)_2_SO_4_, 10 g tryptone, and 5 g yeast extract per liter of water supplemented to a final concentration of 50 mM Tris (pH 7.4), 0.2 mM CaCl_2_, and 10 mM MgSO_4_). Flasks were incubated at 37°C while shaking at 120 rpm. Every 24 hours, 100 μL from each community was transferred into new flasks containing 10 mL of fresh media. Populations were propagated for 21 days. Each day, we measured phage densities by centrifuging cultures for 5 minutes at 3900 x *g* to pellet cells, serially diluting the supernatant, and spotting 2 μL aliquots onto infused soft agar lawns (LB agar except with 0.7% w/w agar and inoculated with ∼10^8^ cells WT). Every 3 days, aliquots were preserved by freezing at –70°C in 15% v/v glycerol.

Φ21 uses LamB as its native receptor. To monitor the coevolution experiment for phage receptor use innovation, we also aliquoted 2 μL of undiluted supernatant onto lawns of KEIO strain JW3996 (*ΔlamB*) each day. When phages demonstrated the ability to lyse *ΔlamB* cells, we spotted them onto a suite of knockout hosts missing LamB plus one additional outer membrane protein in order to determine the novel receptor^31^. Phages could not lyse ΔLamB ΔOmpC cells, revealing that OmpC was the new receptor. We began to test coevolving phages on lawns of ΔLamB ΔOmpC cells, allowing us to discover another iteration of receptor use innovation. We determined that the third receptor was OmpF by first sequencing whole genomes of coevolved resistant bacteria isolated from the experiment, revealing mutations in *ompF*, and then spotting phage supernatant on engineered ΔLamB ΔOmpC ΔOmpF hosts. The inability of phages to infect ΔLamB ΔOmpC ΔOmpF hosts confirmed OmpF as the final receptor.

### Strain isolation and culture techniques

To isolate bacteria, scrapes (∼2 μL) of frozen preserved communities were streaked onto LB agar plates and incubated overnight at 37°C. Then, colonies were picked randomly and streaked twice more to obtain clonal strains devoid of phage. Finally, purified strains were grown overnight at 37°C and preserved by freezing, as described above. Bacteria were named by isolation day and a unique identifier (e.g., T3_1 is day 3 isolate 1).

To isolate phages, scrapes were first suspended in 1 mL Tris-LB. Then, aliquots were inoculated into molten (∼55°C) soft agar infused with cells, poured over LB agar plates, and incubated overnight at 37°C. We isolated ∼12 phages per timepoint. Three phages were isolated on soft agar infused with WT and, to enhance diversity of receptor use types, we also isolated 3 phages on ΔOmpC ΔOmpF, ΔLamB ΔOmpF, and ΔLamB ΔOmpC hosts. From some timepoints, we were unable to isolate phages on particular hosts/receptors. For example, on day 3, we could not recover phages that used OmpF (ΔLamB ΔOmpC hosts). To purify phage isolates, plaques were randomly picked and re-plated with respective isolation hosts twice more. Finally, purified plaques were picked into Tris-LB with respective isolation hosts and grown overnight at 37°C. The next day, phages were filtered through 0.2 μm filters to remove cells and preserved by freezing, as described above. Phages were named by isolation day, isolation host (WT, C-F-, L-F-, L-C-), and a unique identifier (e.g., T03_C-F-_1 was isolated from day 3 on ΔOmpC ΔOmpF hosts and was the first strain picked).

### PBIN measurements

We measured phage-bacteria interaction networks (PBINs) by testing pairwise infections between phages and bacteria in conventional spot assays. Bacterial freezer stocks were inoculated into 18 mm tubes containing 4 mL of LB media. Phage freezer stocks were inoculated into 4 mL of Tris-LB with ∼10^7^ cells. All tubes were incubated overnight at 37°C shaking at 220 rpm. For measuring the full PBIN (Fig. 1a), phages were grown with respective isolation hosts. However, we later found that results were more robust when phages were grown with WT (hosts/receptors used for isolation were not always preferred by phages, leading to large differences in phage titers). Thus, proceeding PBINs were measured with phages grown with WT (Extended Data Fig. 3a, b).

To conduct spot assays, bacteria (∼10^8^ cells) were infused into molten soft agar and poured over LB agar plates. Overnight phage cultures were centrifuged for 10 minutes at 3900 x *g* to pellet cells. Then, 2 μL aliquots of undiluted phage supernatant were spotted onto bacteria-infused soft agar plates and incubated overnight at 37°C. The next day, plates were visualized. Phages that produced visible zones of cell lysis were deemed able to infect and phages failing to produce any visible clearing were deemed unable to infect.

### Network analyses and statistics

We used the BiMat MATLAB library to conduct bipartite network analyses (see documentation at https://bimat.github.io)^28^. The number of modules in the network was optimized by maximizing the modularity metric *Q*_b_. Initial community detection was calculated using the LP-BRIM algorithm^30^. To test the multi-scale nature of patterns we calculated and tested nestedness and modularity for module 1 and module 2 & 3 where nestedness was based on the NODF algorithm^29^ and modularity was based on the LP-BRIM algorithm^30^. Finally, we calculated and tested nestedness of each module using NODF. All results were statistically tested against a null-model where the overall and marginal connectances are held fixed on average. We performed 1000 runs for significance. All other statistical tests were conducted in R (version 4.1.1)^42^. All code and data for network analyses are available at https://github.com/aluciasanz/nestedness_modularity_pbin_analysis and for all other analyses and figures at https://github.com/joshborin/phi21multiscalePBIN.

### Characterizing phage receptor use

Phage receptor use was determined by utilizing a suite of receptor knockout hosts (see Strains). Each host presents one of three relevant receptors, allowing us to pinpoint which receptor(s) each phage could use. To estimate the frequency of receptor use in whole population samples (Fig. 1a), scrapes of frozen preserved communities from various timepoints were suspended in Tris-LB, inoculated into soft agar infused with WT or knockout hosts, and poured over LB agar plates. After incubating plates overnight at 37°C, we counted the number of plaques on each host and calculated the frequency compared to WT.

To determine the receptor use of individual phage genotypes (Fig. 2b, Extended Data Fig. 7), phage freezer stocks were scraped into Tris-LB with WT and incubated overnight at 37°C. Then, cells were pelleted by centrifugation at 3900 x *g* and 2 μL aliquots were spotted onto lawns containing WT cells or knockout hosts. Plates were incubated overnight at 37°C and phages were deemed able to use a given receptor if they could lyse respective knockout hosts.

### Bacteria and phage genomics

Genomes were extracted and analyzed as described in Borin et al., 2021^43^, except with the following differences: Bacteria were grown in LB. Phages were grown with WT in Tris-LB. Extracted genomes were sent to SeqCenter (Pittsburgh, PA) where they were indexed and sequenced on an Illumina NextSeq 2000. To conduct whole population sequencing (Fig. 2a), genomes were extracted as described for bacteria, except overnight cultures were inoculated with scrapes from frozen preserved communities instead of purified bacterial isolates. After sequencing, whole populations were analyzed using the computational analysis pipeline *breseq* (version 0.35.0) set to polymorphism mode^44^.

### Genetic engineering

To engineer phages and bacteria, we used a modified protocol^45^ for Multiplexed Automated Genome Engineering (MAGE)^46^, which employs the Lambda Red recombineering system (residing on plasmid pKD46) to recombine double-stranded DNA fragments or 90 bp single-stranded oligonucleotides (oligos) to edit genomes (Table 1). To make the wildtype Φ21 strain lytic, we conducted MAGE on K-12 lysogens using an oligo that introduces two successive stop codons and a 1 bp deletion frameshift early in the *cI* gene. After electroporation, ∼10^8^ WT cells were added to recovery cultures as fodder to enrich for successfully engineered lytic phages. Recovery cultures were then plated on lawns of WT and clear plaques, indicative of lytic mutations, were isolated, confirmed by whole genome sequencing, and preserved by freezing as described above.

**Table 1.**
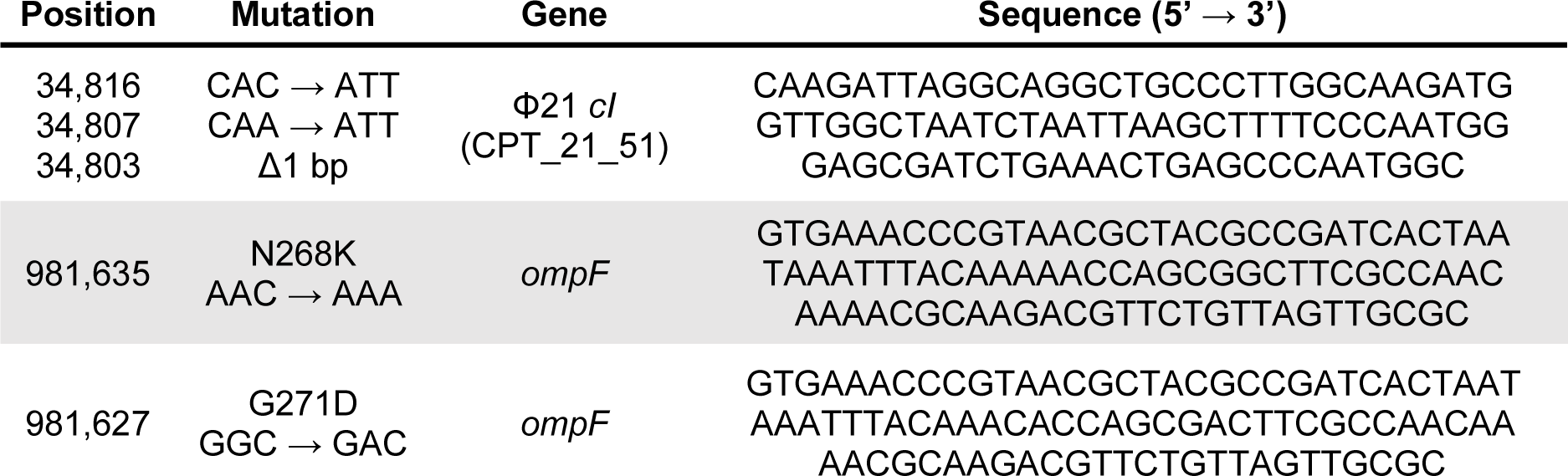
MAGE oligos used in the study.

To engineer bacteria, we used MAGE to introduce *ompF* mutations into ΔLamB ΔOmpC hosts. Mutations observed in module “entry-level” strains (S53R, E93K and G347D) were engineered using mutant *ompF* gene fragments instead of oligos. Fragments were obtained using PCR to amplify *ompF* from coevolved bacteria containing mutations of interest (T12_12, T15_14) (Primers in Table 2). Then, MAGE was used to recombine mutated *ompF* fragments into ΔLamB ΔOmpC hosts. Strains that were successfully engineered with the S53R mutation were used as a scaffold to introduce additional *ompF* mutations using oligos (Table 1).

**Table 2.**
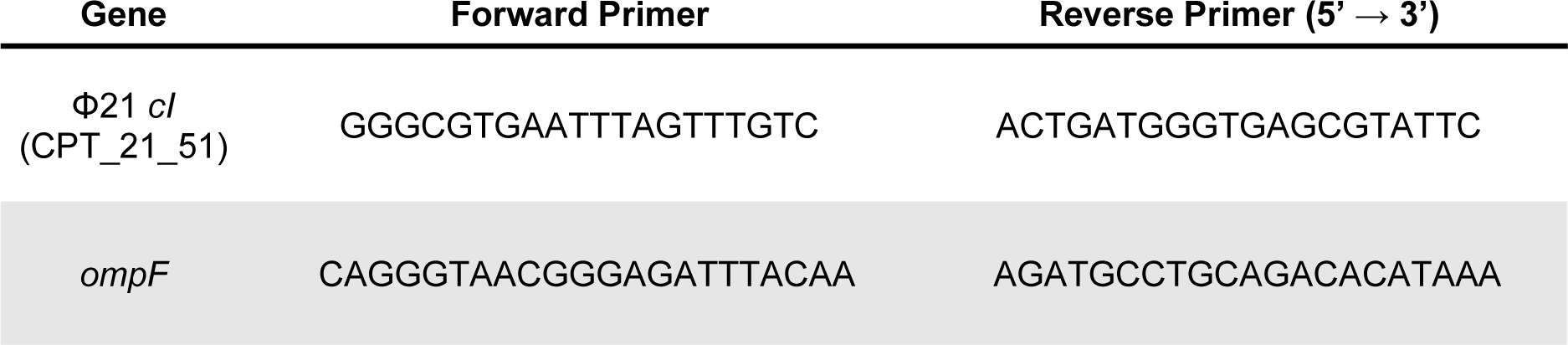
Primers used in the study.

After electroporating ΔLamB ΔOmpC hosts with either *ompF* PCR fragments or MAGE oligos, cells were recovered for 2 hours at 37°C and then 40 μL was transferred into 4 mL of fresh LB media and incubated for another 4 hours. This allowed transformants to divide, thus diluting away OmpF proteins that were expressed before electroporation. Then, we enriched for successful transformants using phage selection in 96-well plate reader experiments. Recovered cultures were serially diluted and 10 μL aliquots were added to wells containing 150 μL of Tris-LB and 50 μL of phage lysate. Plates were incubated at 37°C and after ∼6 hours. When optical density increased above the limit of detection (OD600 ≈ 0.15), suggesting growth of resistant transformants, bacteria were streaked from cultures onto LB agar plates for isolation, as described above. For phage selection, we used the PBIN to choose phages that could infect the pre-engineered form of OmpF but not the successfully transformed variant of the OmpF. After purifying bacterial isolates, we confirmed that engineering was successful; PCR was used to amplify *ompF* fragments (primers in Table 2) which were sent to Azenta Life Sciences (La Jolla, CA) for Sanger sequencing.

### Bacterial fitness competitions

Competitions were conducted between query strains (coevolved bacterial isolates or K-12 WT, which are *manXYZ*^*+*^) and a *manXYZ*^*–*^*-*marked common competitor (bacterial isolate T06_6). To differentiate strains, tetrazolium-mannose agar (Tet-man) indicator plates (10 g tryptone, 1 g yeast extract, 5 g NaCl, 16 g agar, 10 g mannose per liter of water, and supplemented to a final concentration of 0.005% triphenyl tetrazolium chloride [TTC] indicator dye) were used for plating. Bacteria were grown from freezer stocks overnight, as described above. The next day, query strains and the marked competitor were inoculated into flasks containing 10 mL Tris-LB in a 1:1 or 99:1 ratio to a final volume of 100 μL. Upon inoculation, flasks were mixed and aliquots were diluted and plated to enumerate initial densities (T_0_). After incubating for 24 hours at 37°C, competitions were again diluted and plated to obtain final densities (T_F_). Finally, relative fitness (W) was calculated for each strain where W = M_A_ / M_B_ and where M_A_ = ln(T_F_ / T_0_) of the query strain and M_B_ = ln(T_F_ / T_0_) of the marked competitor. To relate coevolved isolate fitness values to their common WT ancestor, we divided all fitness values by relative fitness of WT to the marked competitor.

### Protein structures

To visualize the positions of mutations responsible for module formation and within-module nestedness, we modeled the protein structures of host receptor OmpF and phage host recognition protein J. For OmpF, we used a structure solved by x-ray diffraction (PDB 3HW9, 2.61 Å resolution) and for J we used a publicly available version (v1.5.2)^47^ of AlphaFold^48,49^ to predict protein structures. We relied on predicted J structures because no experimentally solved structures are available for Φ21 J or closely related proteins. For example, λ phage J is unsolved, however previous work on predicted structures of λ’s J shows that well-characterized mutations occur in expected regions of the protein: mutations affecting host tropism occur on finger-like projections on the outward surface of the protein and mutations affecting thermostability are located distal from the surface^50^. We modeled a truncated version of J containing the 153 most C-terminal amino acids (positions 1008-1160, as in Strobel et al. (2022)). Protein data bank files (.pdb) were then visualized and annotated in ChimeraX (v1.5)^51^.

### Replay coevolution experiment

To study the repeatability of dynamics reported in our study, we conducted 9 additional replicate populations as described above but for 24 days. Because we expected phages to evolve new receptor usage, we measured the titer of phage populations on each of our knockout hosts, allowing us to capture the population dynamics of phage receptor innovation with higher resolution (Extended Data Fig. 5, bottom).

