## Extended Data for "Rapid bacteria-phage coevolution drives the emergence of multi-scale networks"

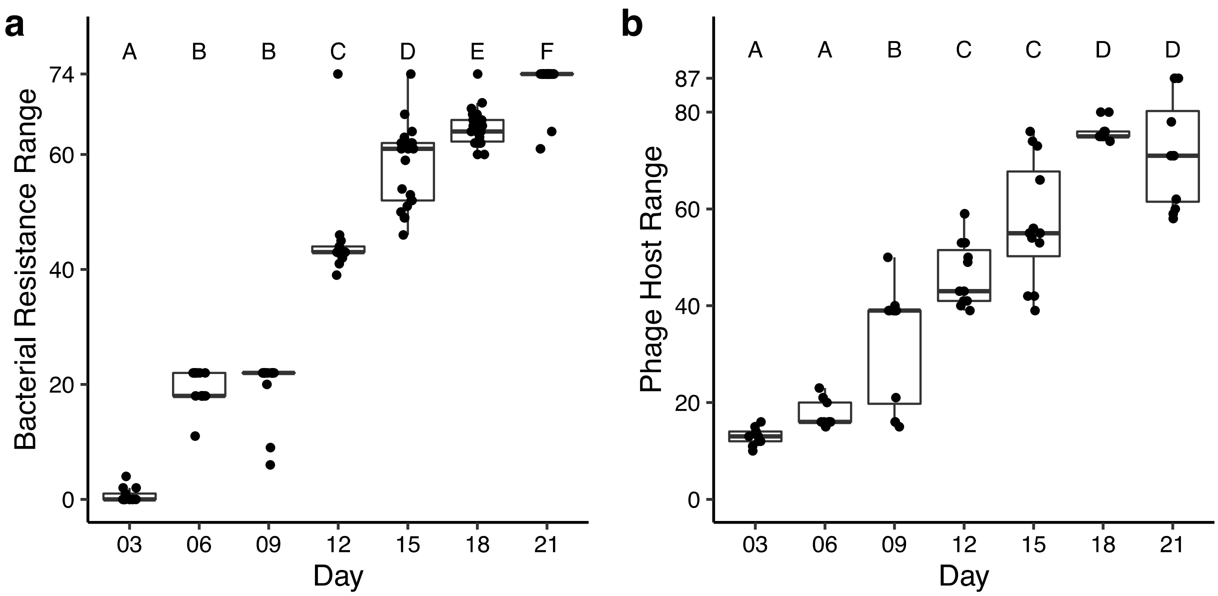


**Extended Data Fig. 1. Bacterial resistance and phage host range increase during coevolution. a,** Resistance range of bacteria isolated from various days of the coevolution experiment, calculated as the number of phage isolates each bacterial isolate was resistant to. **b,** Host range of phages isolates from various days of the coevolution experiment, calculated as the number of bacterial isolates each phage could infect. Infectivity or resistance was determined by the presence or absence of clearing via spot assay. Increase in resistance (**a**) and host range (**b**) over time was statistically significant (linear model, $p<0.001$). Capital letters at top distinguish significantly different groups (ANOVA with post hoc Tukey's HSD $p<0.01$).

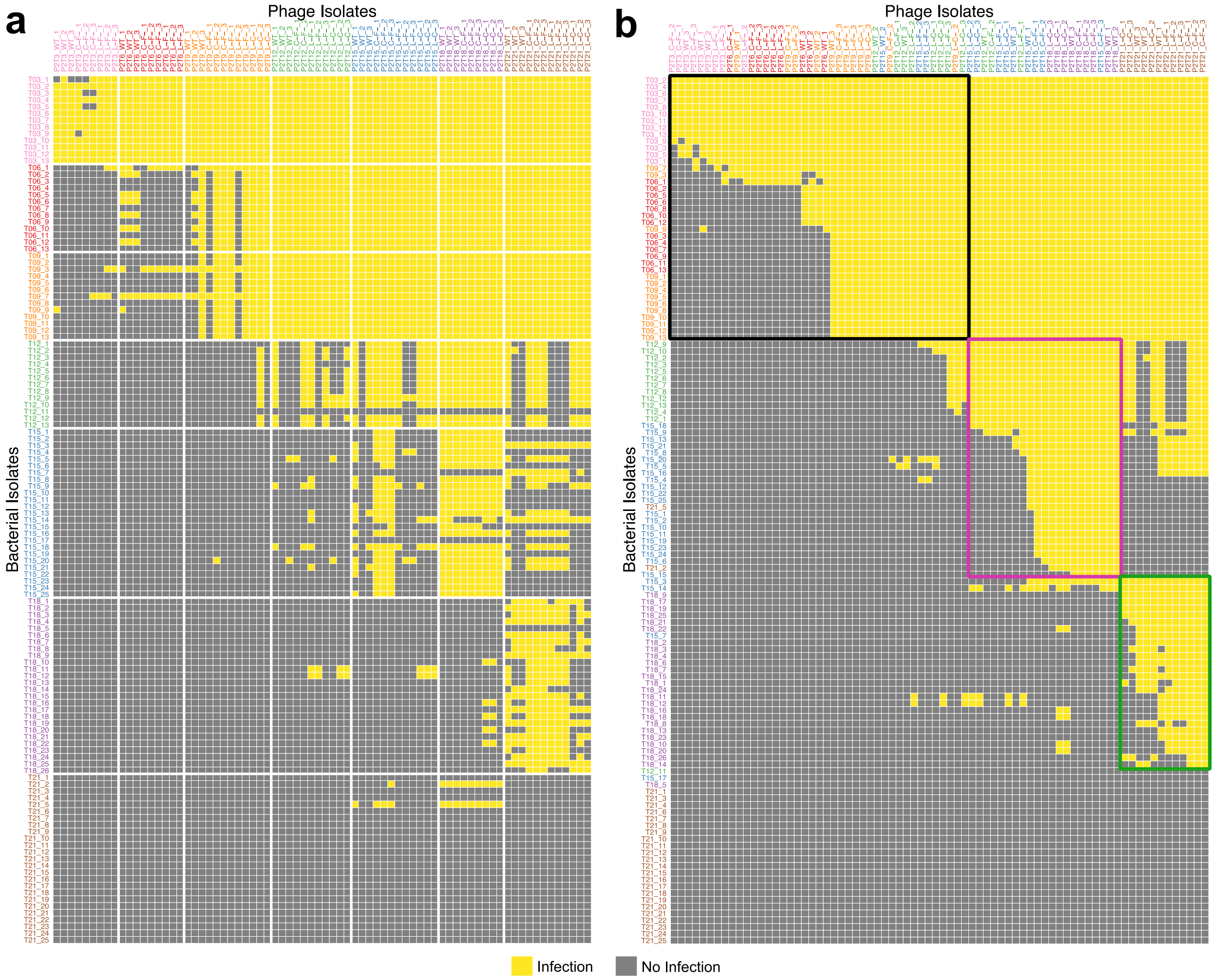


**Extended Data Fig. 2. Full phage-bacteria interaction network (PBIN) including isolate labels. a,** Phage-bacteria interaction network (PBIN) of 9,472 pairwise infections between *E. coli* and Φ21 isolates from various days of coevolution. Isolate labels colored by day of isolation (e.g., day 3, T03, pink). **b,** Reordered PBIN using time-agnostic BRIM algorithm. Modules 1, 2, and 3 are indicated in black, pink, green, respectively. Data also presented in main text Fig. 1a and b.

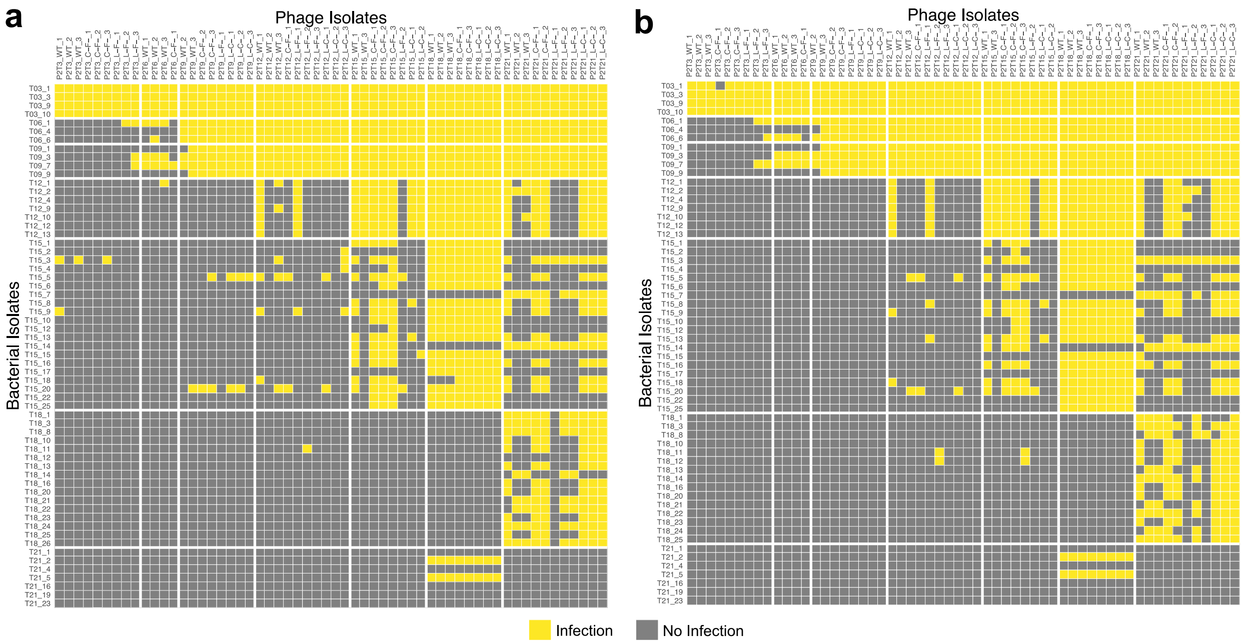


**Extended Data Fig. 3. Independent PBIN measurements are highly repeatable. a,** Time-ordered phage-bacteria interaction network (PBIN) comprised of 3,654 pairwise infections. Isolates in the PBIN are a subset of *E. coli* and Φ21 isolates from the full PBIN. Isolates in the full PBIN were omitted from remeasurement if they had redundant resistance or infectivity profiles within the same timepoint. **b,** Independent measurement of the PBIN in **a**. Both PBINs are significantly correlated with the full PBIN (Fig. 1a; p=0.001, r=0.83, Mantel test).

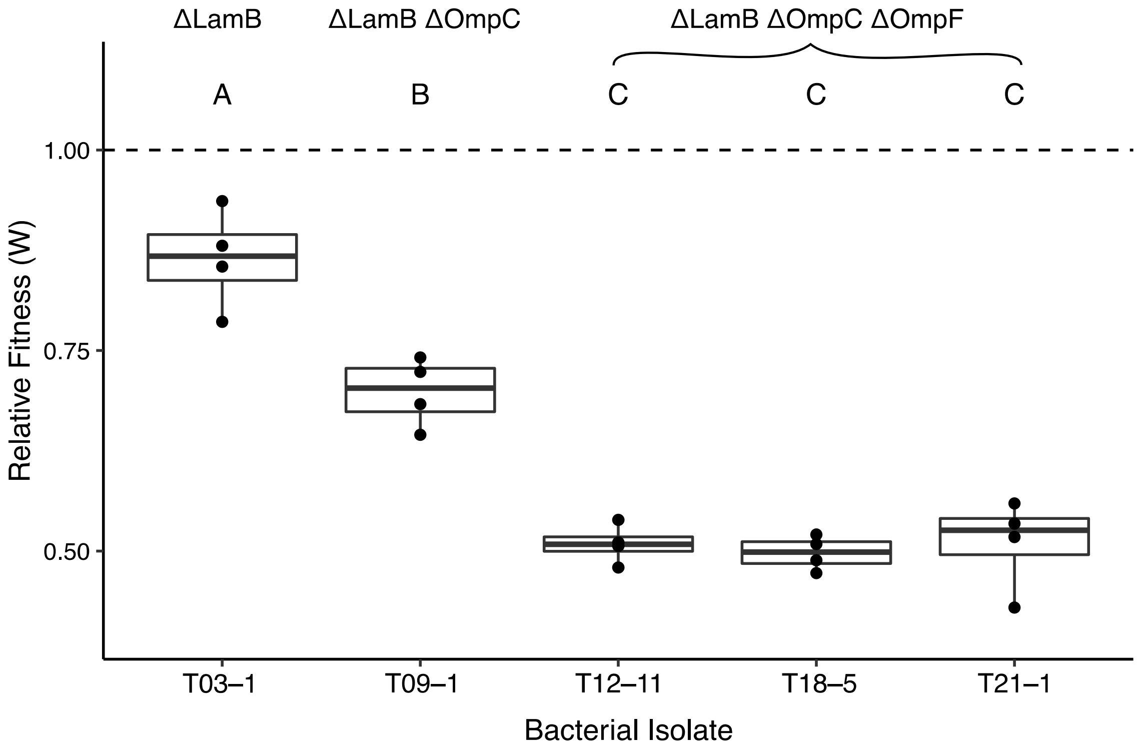


**Extended Data Fig. 4. Fitness costs associated with the sequential removal of host receptors.** Fitness of coevolved bacterial isolates relative to common ancestor K-12 WT (dashed line), measured in quadruplicate. Isolates are labeled and ordered by day of isolation (e.g., T03, day 3). Letters (A, B, C) distinguish statistically different groups (Tukey's HSD, $p<0.05$). Loss of receptor function, due to nonsense mutations or large deletions, are annotated at top.

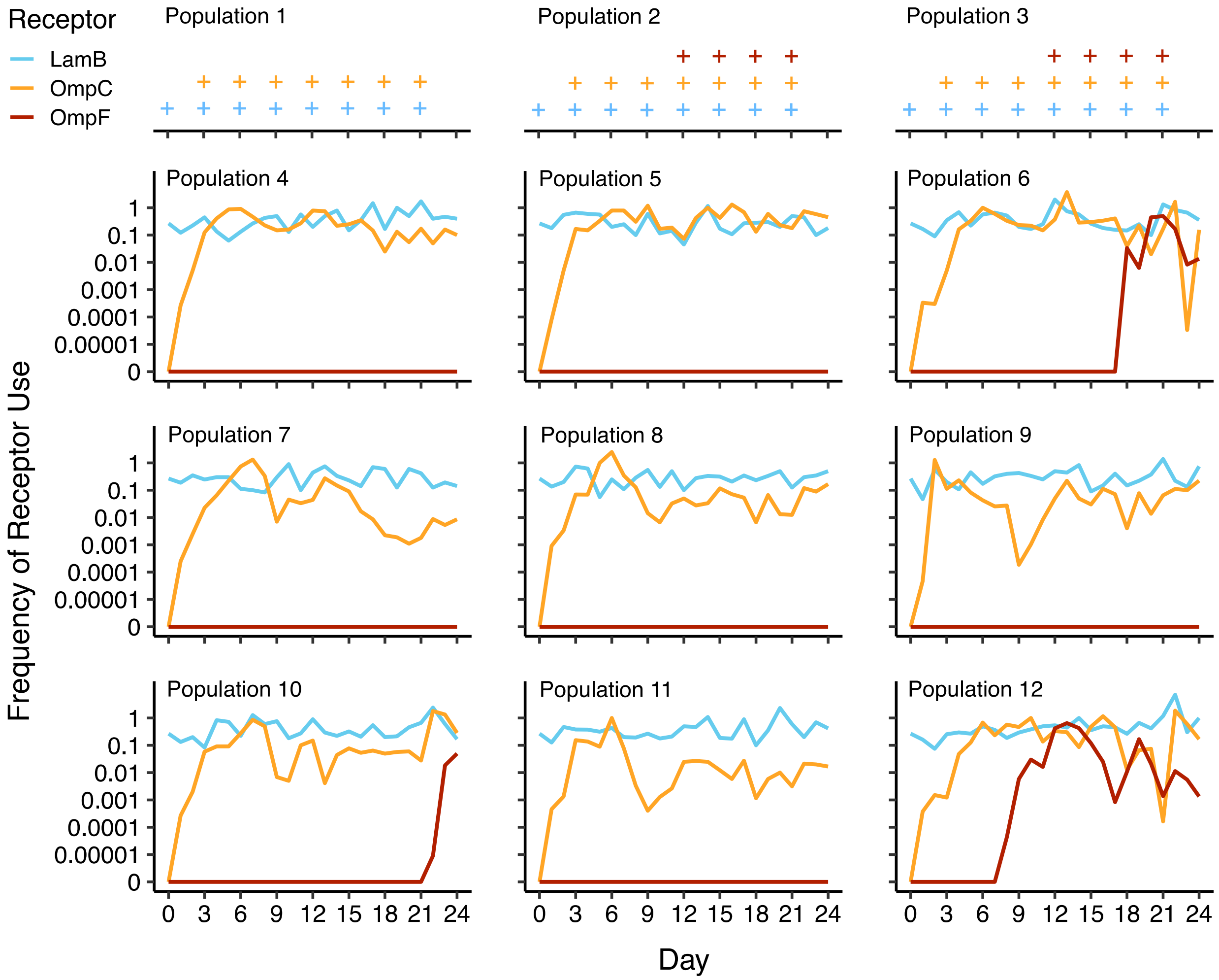


**Extended Data Fig. 5. Independently coevolved phage populations show repeatable dynamics.** For populations 1–3, coevolved for 21 days in the first trial, symbols (+) indicate days where phages in the population could infect through LamB (blue), OmpC (orange), or OmpF (red), determined by plating phages on agar infused with ΔOmpC ΔOmpF, ΔLamB ΔOmpF, or ΔLamB ΔOmpC K-12 hosts. PBINs in the main text were derived from population 2. For populations 4–12, coevolved for 24 days in the second trial, we anticipated receptor innovation so we measured phage titers on each receptor by plating on agar infused with ΔOmpC ΔOmpF, ΔLamB ΔOmpF, ΔLamB ΔOmpC, or WT K-12 hosts.
